# Genetic context drives evolution of divergent antibiotic survival phenotypes in *Staphylococcus epidermidis*

**DOI:** 10.64898/2026.05.29.728712

**Authors:** Carolin M. Kobras, Carlota Losa-Mediavilla, James C. Littlefair, Elizabeth A. Cummins, Charlotte Woolley, Seungwon Ko, Priyanshu Singh Raikwar, Keith A. Jolley, Samuel K. Sheppard, Mathew Stracy

**Affiliations:** Sir William Dunn School of Pathology, University of Oxford, Oxford, United Kingdom; Institute of Microbiology and Infection, Department of Microbes, Infection and Microbiomes, College of Medicine and Health, University of Birmingham, Birmingham, United Kingdom; Ineos Oxford Institute for Antimicrobial Research, Department of Biology, Life and Mind Building, University of Oxford, Oxford, United Kingdom

**Keywords:** antibiotic resistance, antibiotic tolerance, *Staphylococcus*, β-lactam antibiotics, evolution, epistasis, genetic variation, population genomics

## Abstract

Effective treatment of infections is a global challenge, complicated by bacteria’s capacity to endure antibiotics. Survival under drug pressure is often driven by interactions between genetic factors rather than single genes. To investigate the genomics underlying antibiotic survival, we analysed *Staphylococcus epidermidis* isolates from clinical infections and carriage using high-throughput phenotyping, population genomics, and directed evolution. We observed widespread multidrug resistance, with strong links between specific genomic elements and resistance. All isolates harbouring *mecA* were resistant to oxacillin, though minimum inhibitory concentrations varied significantly, suggesting modulation by additional genetic factors. Directed evolution revealed potentiating mutations that enhanced oxacillin resistance in *mecA*^+^ strains. In *mecA^-^*isolates, however, evolution of mutations in the same genes conferred increased survival to oxacillin through antibiotic tolerance. These findings show that antibiotic resistance and tolerance can be genetically connected yet phenotypically distinct, and suggest a complex epistatic genetic landscape that shapes antibiotic survival phenotypes in *S. epidermidis*.

## Introduction

Antimicrobial resistance (AMR) is a major global public health threat^1,2^. With treatment failure continuing to rise, there is an urgent need for coordinated action in surveillance, stewardship, and public health policy. The integration of whole genome sequencing with clinical microbiology has enabled rapid, accurate identification of resistance genes guiding more rapid and precise clinical responses. However, there is increasing evidence that AMR is not only driven by well-known genes but is the result of complex genetic interactions including potentiation, additive effects, and epistasis, that can influence AMR in significant ways. Furthermore, AMR is not the only cause of poor treatment outcomes. Staphylococci employ a range of other survival strategies, including biofilm formation^3^ and antibiotic tolerance^4^, adding further layers of complexity to the genetic landscape underpinning treatment failure. Understanding this genetic complexity is important for developing effective diagnostics, treatments, and public health interventions.

Staphylococci are common skin commensals but can cause infections following invasive procedures or surgery, frequently resistant to antibiotics^5–7^. *Staphylococcus epidermidis* has emerged as a significant nosocomial pathogen, causing infections associated with indwelling devices, including prosthetic heart valve endocarditis and catheter-related bloodstream infections^7–9^. In contrast to *Staphylococcus aureus*, *S. epidermidis* is often seen as a low-virulence, accidental pathogen^10^. However, the rise of hospital-adapted lineages (e.g. ST2, ST5, and ST23)^11^ and discovery of virulence genes^12–14^ linked to biofilm formation, cytotoxicity, and immune activation challenge this view^15^. Furthermore, 75–90% of *S. epidermidis* isolates from nosocomial infections are methicillin-resistant (MRSE)^10,16^, and many exhibit multidrug resistance, further reducing treatment options such as aminoglycosides, fluoroquinolones, rifamycins, and even glycopeptides^9,11,17,18^. As reliance on implanted devices grows, so too does the incidence of *S. epidermidis* infections and the urgency for understanding emergent opportunistic pathogens.

In staphylococci, β-lactam resistance is primarily mediated by the acquisition of *mecA*, which encodes PBP2a, a low-affinity penicillin-binding protein that allows growth in the presence of β-lactams. Located on the mobile genetic element Staphylococcal Cassette Chromosome *mec* (SCC*mec*), *mecA* plays a central role in the emergence and spread of methicillin-resistant staphylococci^19,20^. Expression of β-lactam resistance is further influenced by complex regulatory and genetic interactions involving SCC*mec*-encoded regulators (e.g. *mecI*, *mecR*), β-lactamase regulatory systems (*blaZ, blaI, blaR*), and chromosomal auxiliary factors that modulate cell wall physiology^21,22^. Consequently, *mecA* alone may only confer low-level resistance, suggesting that further genetic factors are required for high minimum inhibitory concentrations (MICs)^23^. In *S. aureus,* mutations in several genes have been recently shown to potentiate β-lactam resistance^23–25,22,26^, typically linked to cellular physiology rather than directly affecting PBP2a or cell wall synthesis. Although the evolution of high β-lactam resistance is becoming increasingly clear in *S. aureus*, it remains understudied in *S. epidermidis*.

Resistance is not the sole reason for treatment failure in staphylococcal infections. These bacteria also use alternative survival strategies such as biofilm formation^27,28^, immune evasion^29–31^, and antibiotic tolerance and persistence^4,32^. Antibiotic tolerance, a phenomenon distinct from resistance but potentially equally concerning in clinical contexts, allows bacterial populations as a whole to better survive the antibiotic challenge, without significant changes in the MIC^33^. Unlike AMR, antibiotic tolerance is not detected by standard clinical susceptibility testing, which is largely based on growth assays. While tolerance is a highly evolvable heritable trait, observed *in vitro*^34–36^ and in infections^37–39^, our understanding of it lags far behind that of resistance.

Here, we used a combination of high-throughput phenotyping, population genomics, and directed evolution to investigate the genomic basis of antibiotic resistance in clinical and carriage *S. epidermidis* isolates. Specifically, we asked how the presence of major resistance determinants, such as *mecA*, influences phenotypic outcomes. Focusing on oxacillin, we explored how genetic background and evolutionary responses to antibiotic pressure shape survival strategies, such as resistance and tolerance. Our results reveal widespread multidrug resistance and strong associations with resistance genes. We further show that the phenotypic effects of newly acquired mutations depend strongly on the presence of *mecA*, suggesting that epistatic interactions between *mecA* and rapidly evolved mutations drive distinct antibiotic survival strategies in *S. epidermidis* isolates.

## Results

### AMR genotypes correlate with resistance above clinical breakpoints in S. epidermidis

A diverse collection of 88 *S. epidermidis* isolates was examined to understand the prevalence of AMR in natural populations (Supplementary File S1). This collection comprised 37 human infection isolates, primarily sampled from wound and bloodstream infections, and 51 isolates derived from asymptomatic skin and nasal carriage. The isolate collection represented known *S. epidermidis* genetic diversity including isolates from all major sequence types (STs) and clades^13,40,41^ (Fig. S1). The MIC of clinically relevant antibiotics, from 10 different drug classes, was measured for all isolates using microbroth dilutions (Fig. 1, Supplementary File S2). This included oxacillin, used for treatment of methicillin-sensitive *S. epidermidis* (MSSE) infections, and vancomycin, used as first-line treatment of MRSE^9,18^. Of the isolates tested, 68% were resistant to at least one antibiotic class, with MICs above the clinical breakpoint and >30% were multidrug resistant, (resistant to three or more antibiotic families)^42^.

**Figure 1:**
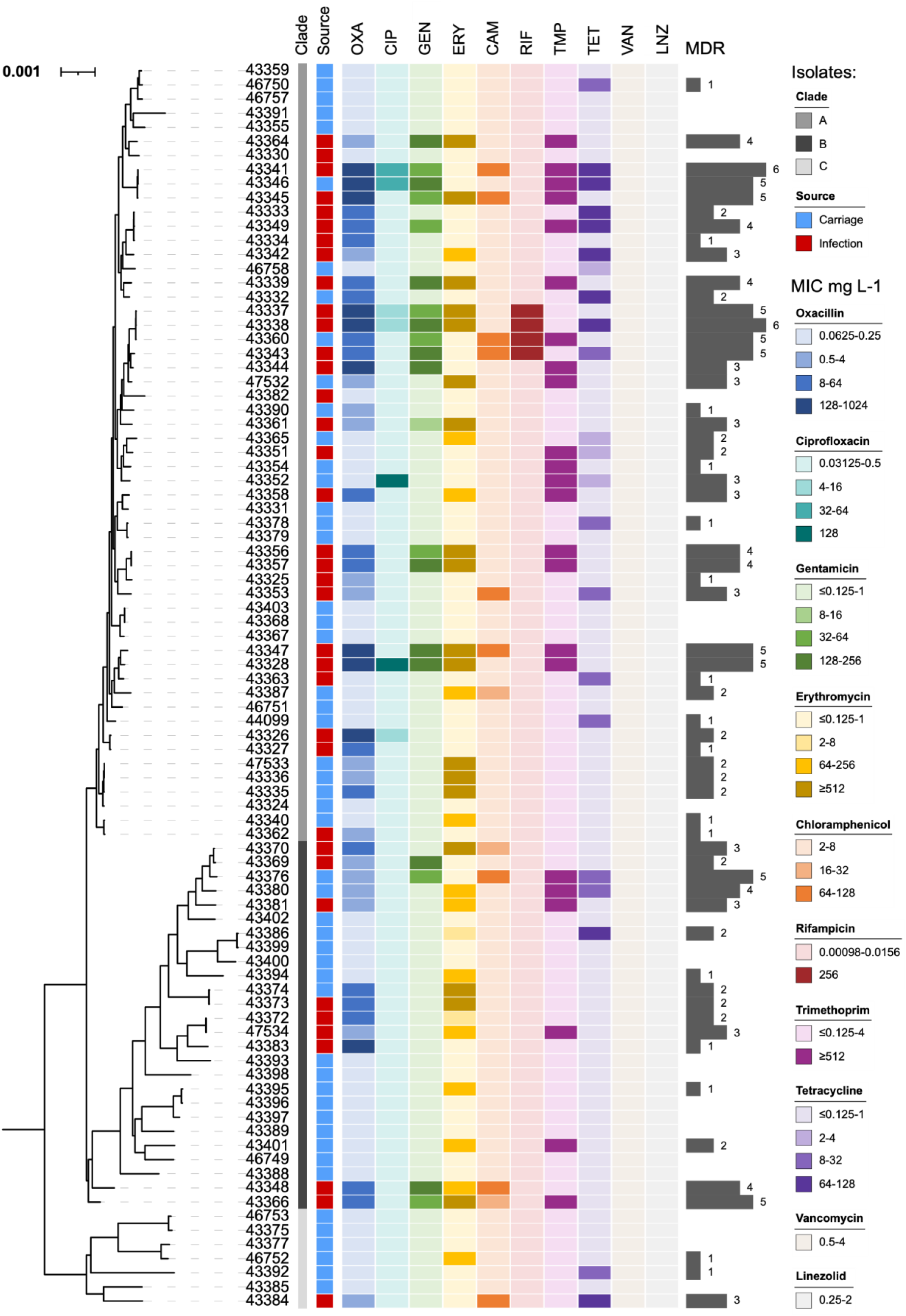
Phylogenetic distribution of phenotypic antibiotic resistance profiles reveal multidrug resistant *S. epidermidis* clusters. Minimal inhibitory concentrations (MICs, in mg L^-1^) of 88 *S. epidermidis* isolates for 10 antibiotic compounds from different drug classes are displayed alongside their phylogeny. The tree scale represents the number of substitutions per site. Different shades of grey and red or blue indicating placement in deep branching clades A-C, and isolation from infections or carriage, respectively. MICs are displayed with the lightest shade indicating susceptibility and darker shades corresponding to increasing levels of resistance. Clinical breakpoints for resistance are (in mg L^-1^): Oxacillin (OXA, dark blue) > 0.25, ciprofloxacin (CIP, teal) > 2, gentamicin (GEN, green) > 2, erythromycin (ERY, yellow) > 1, chloramphenicol (CHL, orange) > 8, rifampicin (RIF, dark red) > 0.06, trimethoprim (TMP, pink) > 4, tetracycline (TET, purple) > 1, vancomycin (VAN, brown) > 4, linezolid (LNZ, light grey) > 4. Multidrug resistance (MDR) is summarised through length of grey bars.

AMR phenotypes can often be inferred from the presence of known resistance genes^43,44^. All isolate genomes were screened for commonly acquired resistance genes and chromosomal mutations linked to AMR across the eight antibiotic classes to which resistance was observed. For many isolates we identified multiple putative genetic resistance determinants for a given antibiotic (Fig. S2, Supplementary File S3). However, in most cases the acquisition of genes or mutation(s) in a single gene was sufficient to predict resistance above the clinical breakpoint (Fig. 2). All oxacillin-resistant isolates harboured *mecA* (PBP2a), all ciprofloxacin-resistant isolates had DNA topoisomerase IV and gyrase substitutions (ParC^S80X^, GyrA^S84F^), and rifampicin resistance was linked to the RpoB^D471E/I527M^ double substitution in the RNA polymerase, while no susceptible isolates carried these resistance determinants. For some antibiotics, resistance was not explained by a single genetic determinant across all isolates. For example, tetracycline resistance was mainly linked to acquisition of *tetK*, *tetL*, or *tetM*. Again, this suggests that a few key genotypes can reliably predict resistance above clinical breakpoints for most antibiotics tested. For some antibiotics, the identified genetic determinants were further strongly correlated with the exact MIC. For example, RpoB^D471E/I527M^ substitutions were always associated with a rifampicin MIC of 256 μg ml^-1^, an over 16,000-fold increase compared to susceptible isolates.

**Figure 2:**
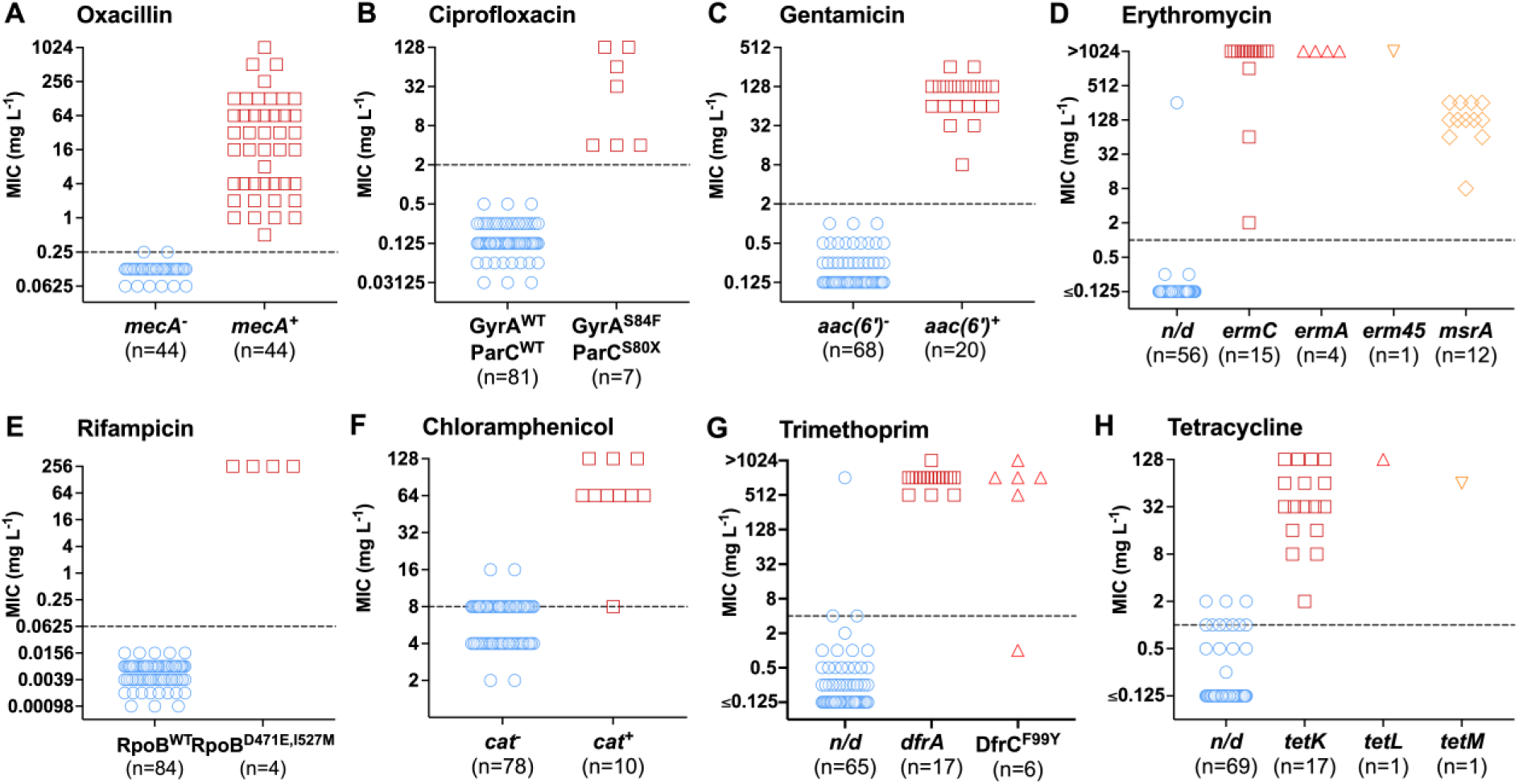
Most phenotypic AMR in *S. epidermidis* isolates can be explained by few resistance determinants. MIC distribution of 88 *S. epidermidis* isolates against A: Oxacillin, B: Ciprofloxacin, C: Gentamicin, D: Erythromycin, E: Rifampicin, F: Chloramphenicol, G: Trimethoprim, H: Tetracycline. Isolates are split into different genotypes based on the most commonly observed genotype that explains resistance. The dashed line indicates the clinical breakpoint for each antibiotic^84^. Dots or squares above this line represent resistant isolates.

### mecA predicts oxacillin resistance but not the minimal inhibitory concentration

For several antibiotics, the gene or mutations which accounted for all resistance above the clinical breakpoint showed poor correlation with the exact MIC. In some cases, the exact MIC could be better explained by incorporating additional known genes or mutations associated with resistance beyond the most predictive individual genetic determinant. For example, while the double substitutions in DNA topoisomerase IV and DNA gyrase (ParC^S80X^, GyrA^S84F^) perfectly correlated with ciprofloxacin resistance, isolates carrying these mutations had MICs ranging from 4 to 128 μg ml^-1^. Isolates with high level resistance (≥ 32 μg ml^-1^) had all acquired additional non-synonymous substitutions in ParC (D84X), the primary target of fluoroquinolones in staphylococci^45,46^.

While the presence or absence of *mecA* perfectly correlated with oxacillin resistance or susceptibility, respectively, the exact MIC of resistant isolates varied over 2,000-fold, consistent with previous observations that *mecA* alone does not fully determine the magnitude of β-lactam resistance^47^. However, this variation could not be explained by known resistance genes or mutations (Fig. 2A). Despite detection of the β-lactamase-encoding gene *blaZ* in 69 of 88 *S. epidermidis* isolates (Fig. S2), there was no significant correlation with high-level oxacillin resistance. Next, we examined the SCC*mec* types, of which 15 distinct classes have been described^19,20^, some of which lack or only have truncated regulatory elements such as *mecI* or *mecR* (Fig. S2, Supplementary File S4). We detected an association between SCC*mec* type I and higher MIC; however, this was observed in three closely related isolates (Fig. S2, Supplementary File 5) and was not sufficient to explain the broad variation in MICs seen in strains with other SCC*mec* types. This highlights a gap in our understanding of what drives the large MIC variability of *mecA^+^* isolates in *S. epidermidis.* It also raises questions about why *mecA*^-^ isolates have such a narrow range of MICs and do not appear to acquire low levels of resistance.

### Parallel evolution of β-lactam resistance potentiators in mecA^+^ isolates

The MIC variation in *mecA^+^ S. epidermidis* isolates can potentially be explained by other genetic factors that enhance resistance. In *S. aureus* mutations in genes affecting regulation of gene expression, modulation of cell signalling pathways, and maintenance of protein stability have been shown to potentiate levels of β-lactam resistance in the presence of *mecA*^22–25^. To understand if similar potentiating mutations can explain high levels of oxacillin resistance in *S. epidermidis*, we performed a directed evolution experiment (Fig. 3A).

**Figure 3:**
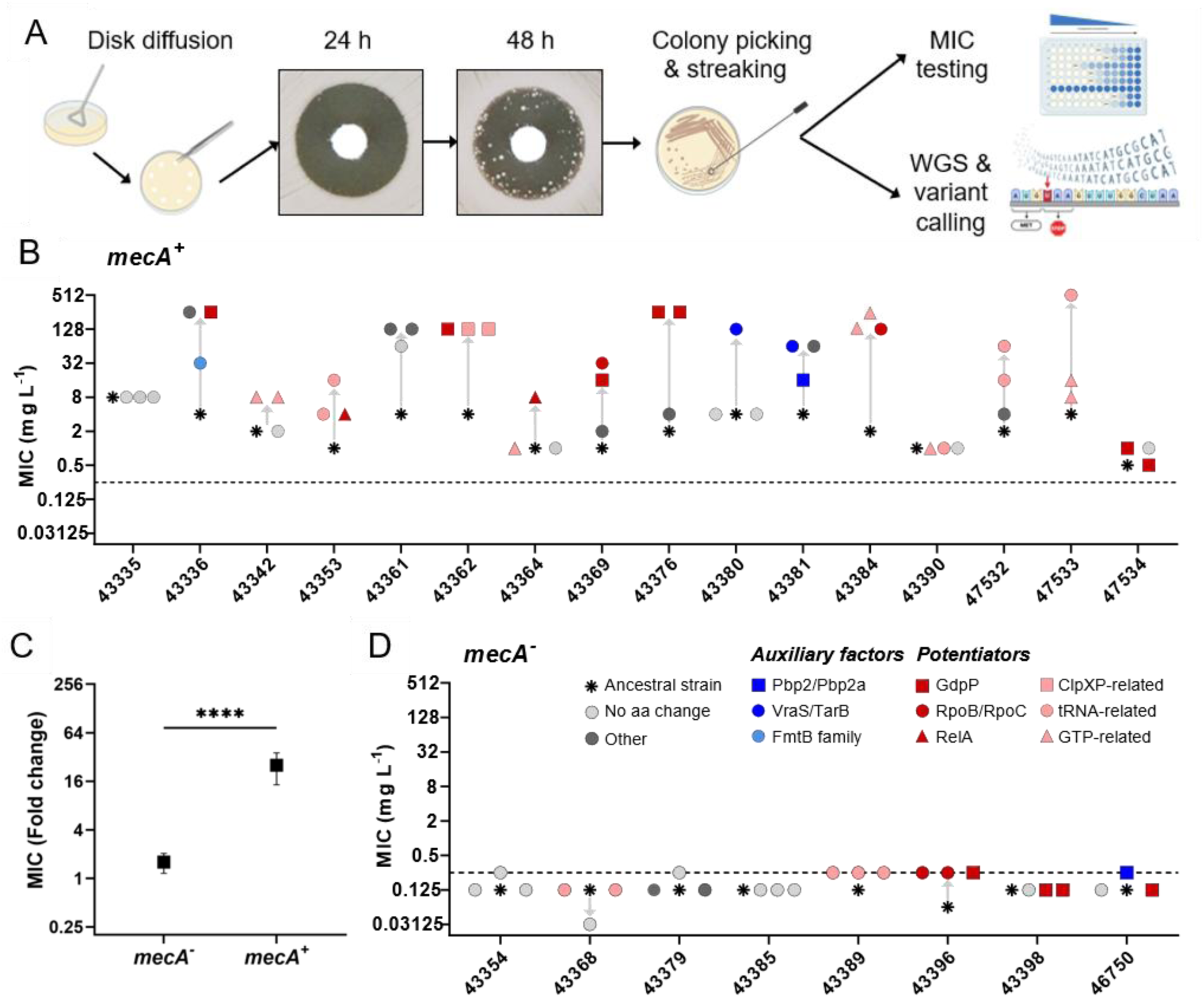
Mutations in genes linked to tolerance can potentiate β-lactam resistance in *mecA*^+^ isolates. A: Schematic of TDtest workflow used to select for oxacillin tolerance and resistance. Created in BioRender. Kobras, C. (2026) https://BioRender.com/3iezucn. B,D: MIC changes of evolved isolates compared to their ancestral strain (n = 3 per ancestral strain) are indicated by changes on the y-axis (grey arrows). MIC changes of evolved isolates are more prominent in *mecA^+^* ancestral isolates (B) than in *mecA^-^* ancestral isolates (D). Shape and colour of symbol indicate changes in genotype: grey square: ancestral strain, grey circle: no amino acid change, red symbols: protein-altering changes in proteins linked to both tolerance and potentiation of β-lactam resistance in *S. aureus*, light red symbols: protein-altering changes in pathways related to tolerance and potentiation, blue symbols: proteins and pathways related to auxiliary factors described for *S. aureus*, dark grey symbols: protein-altering changes in other proteins/pathways. C: Quantification of MIC changes (fold-change) between *mecA^-^* and *mecA^+^* evolved isolates. Statistical significance was assessed by Mann-Whitney test: ****: *p* < 0.0001.

Increased oxacillin resistance readily evolved in most of the 48 *mecA^+^*isolates (36/48, Fig. 3B), with an average 25.3-fold change in oxacillin MIC (Fig. 3C). Interestingly, we observed strong signatures of parallel evolution, with repeated mutations in the same genes across diverse *S. epidermidis* backgrounds. Notably, seven distinct mutations in *gdpP*, a regulator of the second messenger cyclic di-AMP, appeared in *mecA^+^*isolates from five lineages. These included loss-of-function variations, leading to truncated proteins, and missense mutations in proximity to the ligand binding and catalytically active sites, suggesting functional disruption of the protein (Fig. S3). Additional mutations were found in *rpoB*, *rpoC*, *relA*, and *yjbM*, key regulators of transcription^48^ and the stringent response^49,50^. Furthermore, we identified substitutions in genes which had not been associated with resistance before, but elevated β-lactam resistance in our evolution experiment. For example, we detected amino acid changes in YtxJ, a monothiol bacilliredoxin that releases bacillithiols from proteins, which they had acquired as protection from oxidative stress^51^. Notably, we also found two independent amino acid changes mapped to a hypothetical protein predicted to function as an inorganic phosphate transporter, and one in the adjacent regulatory protein (likely PitAR), highlighting repeated non-synonymous changes in genes not previously described as potentiating factors. In addition to potentiator genes, which generally do not seem to be directly involved in cell wall biosynthesis, we also identified alterations in genes involved in the cell wall stress response *(vraS)*, teichoic acid metabolism (*tarB),* cell surface proteins *(fmtB)* and even *mecA* itself. Deletion of these genes have been shown to reduce resistance in *S. aureus*^21,22^, suggesting we here observed gain-of-function mutations. Together, these results demonstrate that increased β-lactam resistance can rapidly evolve in *S. epidermidis*, driven by both known and novel genetic factors beyond *mecA* (Fig. S4, Table S1, Supplementary File S6).

### mecA-dependent evolution of divergent antibiotic survival phenotypes

The MIC of *mecA^-^* isolates in our collection only varied 4-fold (0.0625 to 0.25 μg ml^-1^), compared to over 2000-fold for *mecA^+^*isolates. To investigate why these isolates have such a narrow range of MICs and do not appear to acquire low levels of resistance, we repeated the directed evolution experiment with eight *mecA^-^*, oxacillin susceptible isolates (Fig 3D, dashed line). Of the 24 evolved isolates, none acquired oxacillin resistance above the breakpoint. In fact, most isolates showed only minimal changes in MIC, with an average MIC increase across all isolates of just 1.6-fold compared to 25.3-fold for *mecA^+^* isolates (Fig. 3C). This is consistent with the strict requirement for *mecA* acquisition for breakpoint resistance development in *S. epidermidis* that is described above.

In the evolved *mecA*^-^ isolates, we identified mutations in the same genes in which genetic variation led to increased β-lactam resistance in *mecA^+^* isolates (Table S2, Supplementary File S6). These included *rpoB*, *rpoC*, tRNA ligase genes *lysS* and *thrS,* and the GMP-synthase encoding *guaA*. Strikingly, we identified four independent mutations in *gdpP*, with one leading to an amino acid substitution (T498A) in two distinct genetic backgrounds, again indicating parallel evolution. However, in the absence of *mecA*, these mutations had minimal effect on oxacillin MICs. Instead, time-kill assays revealed that nearly all evolved strains exhibited increased survival under high oxacillin concentrations compared to their ancestral strain, consistent with antibiotic tolerance^33^ (Fig. 4A). The time required to kill 99% of the population (MDK_99_), a proposed standard measure to quantify tolerance^33^, had at least doubled in all but two evolved isolates (Fig. 4).

**Figure 4:**
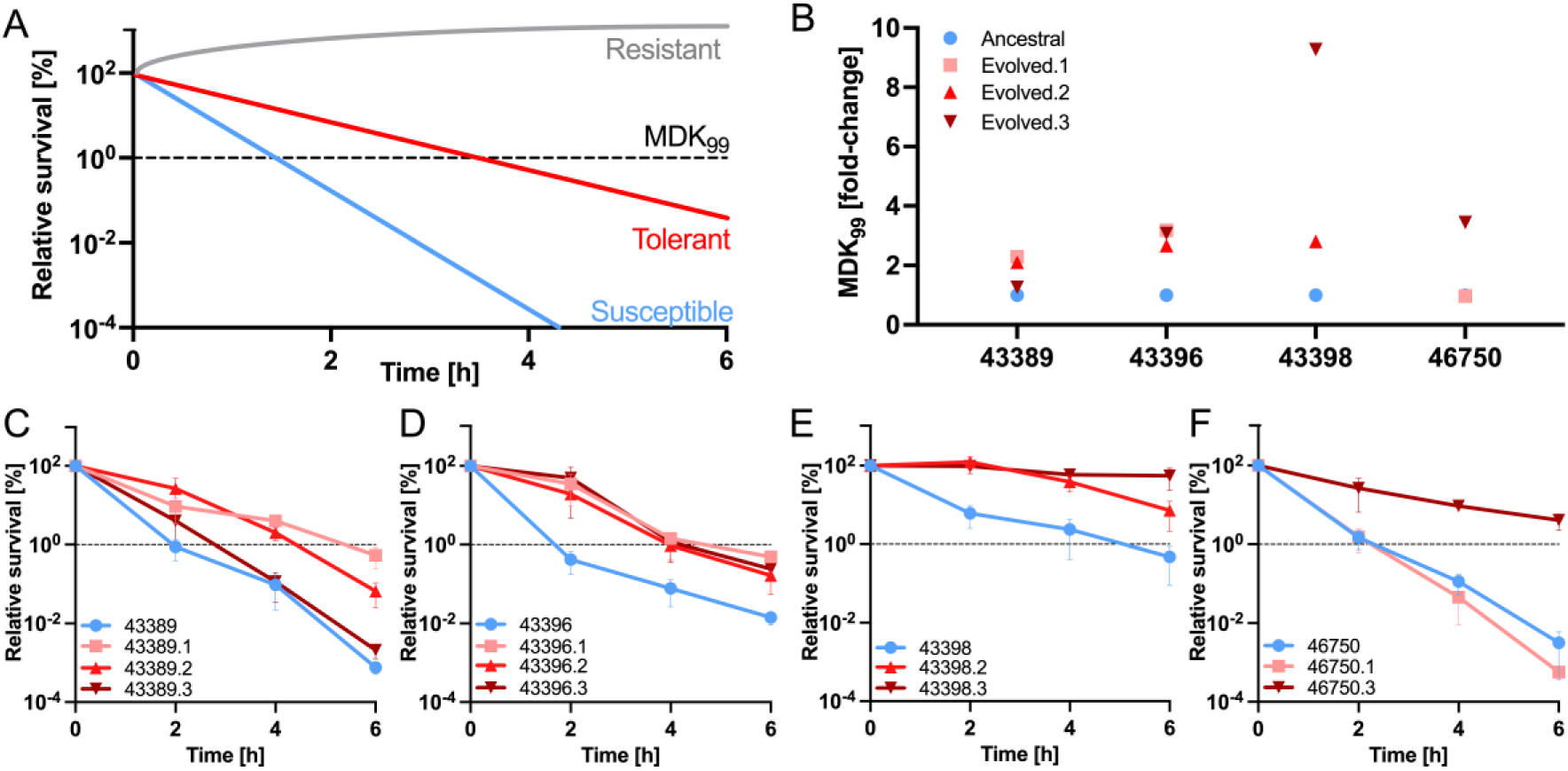
Mutations in potentiating factors increase tolerance in *mecA*^-^ isolates. A: Cartoon showing the different behaviour of resistant, tolerant and susceptible bacteria in time-kill curves. In contrast to resistant bacteria, which can grow in the presence of antibiotics, tolerant bacteria survive the antibiotic challenge better than susceptible ones. The minimum durations for killing 99% (MDK_99_) of bacterial populations (B) and time-kill curves at 100x oxacillin MIC over 6 hours (C-F) are shown for our ancestral strains and all evolved isolates that had amino acid changes. Blue: ancestral strain. Shades of red: evolved strains.

Since killing by β-lactams depends on active growth^52,53^ and growth alterations are a common mechanism of tolerance^33^, we assessed whether tolerance was linked to slower growth in four ancestral strains, alongside two or three respective evolved isolates with non-synonymous mutations. Seven of ten oxacillin-tolerant evolved isolates showed differential growth relative to their ancestors, with five appearing to grow more slowly or with longer lag phase (Fig. S5). In *S. aureus*, *gdpP* and *rpo* mutations have been linked to high ppGpp levels and activation of the stringent response^54–56^. This global regulatory network is associated with stress adaptation, reduced growth, and increased antibiotic survival, suggesting that similar mechanisms may contribute to β-lactam tolerance in *S. epidermidis*^49,50^. Interestingly, evolved isolates of strain 43389, which all carry mutations in tRNA-ligases, showed little or no change in growth (Fig. S5A), despite increased antibiotic survival for 43389.1 and 43389.2. Furthermore, one evolved isolate with a mutation in the penicillin binding protein PBP2, a key enzyme in the cell wall biosynthesis, did not show increased survival despite slower growth, suggesting that not all tolerance may be linked to changes in growth behaviour, and that antibiotic survival strategies in *mecA^-^* isolates may not be limited to tolerance.

Across both evolution datasets (Fig. 3), we observed a significant overlap of genes in which mutations led to either potentiation of oxacillin resistance or tolerance, dependent on the presence or absence of *mecA.* This highlights the importance of the genetic context in which these mutations evolve in determining the resulting survival phenotype, and suggests potential epistatic effects between these tolerance/potentiating mutations and *mecA*.

### Genotypes from in vitro experiment can be identified in natural populations

To contextualise our directed evolution findings in natural bacterial populations^57^, we compared genes that had at least two independent non-synonymous mutations in our experiment to the genetic variation in the genomes of 1,000 *S. epidermidis* isolates (Fig. S6). Despite the vast potential mutational space, both within and across genes that are implicated in potentiating oxacillin resistance, we identified several genes with identical residue changes in the natural population, including GdpP R540 (n=540), CodY N160 (n=4), and A57 in the hypothetical inorganic phosphate transporter (n=1). The recurrence of mutations at the same residues suggests that similar evolutionary trajectories occur in natural populations, highlighting its relevance to real world settings. Of note, we detected an in-frame insertion in GdpP (V323_insRARV) in the genome of a clinical isolate, which resulted in the same alteration to the protein as observed for a *mecA^-^*evolved isolate with increased antibiotic tolerance. This raises the possibility that the insertion may have aided the clinical isolate from a catheter-related invasive infection in surviving antibiotic treatment and thus supports the clinical relevance of our findings.

## Discussion

Treatment failure of *S. epidermidis* infections is an emerging challenge, particularly in healthcare settings where the pathogen is frequently associated with infections involving prosthetic joints and indwelling medical devices^7^. Clinical management relies heavily on antibiotic therapy, particularly the use of β-lactam antibiotics such as oxacillin. However, *S. epidermidis* employs diverse survival strategies to overcome antibiotic exposure, with the development of extensive AMR being among the most prominent^11,58^. Our findings confirm an alarmingly high prevalence of multidrug resistance among a genetically diverse collection of clinical and carriage *S. epidermidis* isolates. Beyond AMR, treatment failure can also be a consequence of a different survival mode: antibiotic tolerance. The rapid emergence of tolerance mutations, followed by the emergence of resistance, has been observed in *S*. *aureus* bloodstream infections^37^. However, in *S. epidermidis*, tolerance remains poorly understood.

Accurate genotype-based prediction of AMR phenotypes has the potential to significantly enhance infectious disease diagnostics. Such predictive tools enable rapid and targeted treatment strategies to improve patient outcomes, reduce antibiotic misuse, and support public health surveillance of AMR. Our genomic analysis demonstrates that a limited set of well-described resistance genotypes reliably correlate with AMR phenotypes above the clinical breakpoint for most clinically relevant antibiotics tested here. Often, the identified AMR genotypes even exhibited strong correlations with the magnitude of resistance, or the precise MIC. In other cases, e.g. ciprofloxacin resistance, this correlation required accumulation of several genetic resistance determinants, which are well documented in the literature and AMR databases^59–61^. Notably, the presence of *mecA* was a perfect predictor of oxacillin resistance in *S. epidermidis* isolates, with isolates harbouring *mecA* consistently exhibiting resistance above the clinical breakpoint. However, in contrast to other antibiotics, neither known resistance determinants as found in the CARD database^59^ nor additional regulatory features such as the SCC*mec* type could explain the widely varying oxacillin MIC range, emphasising the complexity of resistance mechanisms and highlighting the limitations of current resistance databases^59–61^.

Through directed evolution experiments, we identified mutations that could significantly elevate β-lactam resistance in *mecA*^+^ *S. epidermidis* isolates. Although *mecA* promoter mutations or copy number changes can alter oxacillin MIC, we did not identify any such mutations in evolved isolates. Instead, potentiator mutations were found particularly in enzymes involved in transcription, second messenger regulation, and the stringent response. While such potentiating factors have been described in *S. aureus*^23–25,22,62–64^, they had not previously been observed in coagulase-negative staphylococci such as *S. epidermidis*. This study therefore highlights the potential for the rapid resistance evolution of high-level β-lactam resistance in *S. epidermidis,* with potentially important implications for the treatment of these opportunistic pathogens. In contrast to the wide range of MICs observed for *mecA^+^* strains, isolates lacking *mecA* had an intriguingly narrow MIC range, never acquiring low-level oxacillin resistance above the breakpoint, not even through a directed evolution experiment. Nevertheless, the isolates evolved similar mutations in the same genes as observed for *mecA*^+^ isolates. However, in the absence of *mecA*, these mutations seemed to only cause minimal MIC changes. Instead, most evolved isolates (8/10) tested showed increased survival at high oxacillin concentrations, known as antibiotic tolerance. Antibiotic resistance and tolerance are generally regarded as two distinct survival mechanisms^33^ and, accordingly, the genes underpinning them have been reported separately. Our findings, however, suggest that these survival strategies can be genetically interlinked and highlight a significant overlap of genetic factors involved in both tolerance and high-level oxacillin resistance in *S. epidermidis*.

In fact, several of the resistance potentiating genes we identified here have been associated with antibiotic tolerance in *S. aureus*^4,32^, and some of them, including *rpoB* and *rpoC,* even in *S. epidermidis*^38^. Consistent with this, we identified several potentiating factors associated with the activation of the stringent response, changes in the activity of RNA polymerase, the purine salvage pathway, and the oxidative stress response, previously implicated in tolerance^65,32^. Moreover, increased levels of the bacterial second messenger cyclic-di-AMP, often mediated through mutation of the phosphodiesterase-encoding *gdpP*, have been shown to elevate both β-lactam resistance and tolerance in several Gram-positive bacterial species^66^, including *S. aureus*^67–70,64^, *Enterococcus faecalis*^71^ and *Streptococcus pneumoniae*^72^.

Our findings show that chromosomal mutations in those genes can evolve rapidly under antibiotic selection, but that the resulting phenotypically distinct survival mechanisms (increased tolerance versus resistance), suggesting potential epistatic interactions driven by the presence or absence of *mecA*. This work shows that the genetic background of *S. epidermidis* isolates shapes the phenotypic outcome under antibiotic selective pressure, beyond the previously reported acceleration of subsequent resistance evolution of tolerant bacteria^37,69^. Future studies using isogenic backgrounds will be important to resolve the specific effects of individual mutations and their interactions with *mecA*.

The exact mechanism that drives this phenotypically different survival following evolution of similar mutations remains puzzling. Changes to bacterial growth, such as an extended lag phase or an inherently slower growth rate, are an important cause of antibiotic tolerance^33^. Consistent with this, several evolved isolates displayed growth defects. Mutations in regulators of the stringent response, which result in increased ppGpp levels in the cell, have recently been shown to bypass the requirement for PBP1-mediated septal peptidoglycan synthesis in *S. aureus*^54^. PBP2a mediates β-lactam resistance through its low affinity for these antibiotics, enabling it to functionally compensate for the inhibited PBP2 and sustain peptidoglycan cross-linking^54,73^. Together, this appears to facilitate cell division via an alternative mechanism to the classical septal ring model in the presence of antibiotics and results in high β-lactams resistance.

Interestingly, we observed clear signatures of parallel evolution, with similar mutations recurring in the same genes across genetically diverse *S. epidermidis* isolates. For both tolerance^4,32^ and potentiating factors^22^, it is now known that the respective phenotypes can be conferred by one of multiple mutations across several target genes. Despite the large number of potential mutational targets, the repeated selection of mutations in specific genes suggests that some mutational paths may be favoured, even when alternative routes are theoretically available. Some isolates appeared particularly biased towards evolving along similar mutational routes, including independent occurrences of distinct mutations in *gdpP* or tRNA ligase genes. Although our dataset is not large enough to quantify these patterns robustly, this seemingly non-random distribution hints at constraints or biases in the evolutionary trajectories different isolates can follow and raises questions about how genetic background influences mutation outcome beyond *mecA* interactions, and to what extent does it compensate for fitness costs.

While the extent to which findings from directed evolution experiments in laboratory settings translate to natural environments often remains uncertain, we here identified some mutations affecting the same residues in both our laboratory-evolved strains and natural *S. epidermidis* isolates. For example, we identified a four amino acid insertion in GdpP in a *mecA^+^* clinical isolate that alters the protein in the same way as in one of our *mecA^-^* evolved strains. Given that this insertion was associated with increased antibiotic tolerance *in vitro*, its presence in a *mecA^+^* clinical isolate from an invasive infection raises the possibility that this tolerance-associated mutation may have acted as a resistance potentiator in natural populations. The *mecA* locus has been shown to undergo rapid gene gain and loss events in *S. epidermidis* colonising humans^74^. As tolerance mutations have been observed to facilitate subsequent resistance, both in the lab and in the clinic^37,69,75–77^, it is therefore conceivable that tolerance-conferring mutations may evolve in the absence of *mecA*, and act as potentiators upon *mecA* acquisition, allowing a single-step evolution from oxacillin susceptible to high-level resistant. Alternatively, potentiator mutations in a *mecA^+^* isolate may still allow for increased oxacillin survival if *mecA* is lost. Interestingly, tolerance-conferring mutations, such as *rpoC*, can evolve in response to non*-*β-lactam antibiotics^38^. This raises the possibility that selective pressures from unrelated antibiotics can collaterally potentiate oxacillin MICs, or, alternatively, that potentiation to high oxacillin resistance may indirectly confer tolerance to other antibiotics.

In conclusion, this study reveals widespread multidrug resistance in *S. epidermidis* natural isolates, with strong links between specific genomic elements and resistance. We uncover a complex genetic landscape underlying β-lactam survival in *S. epidermidis*, highlighting distinct yet overlapping roles for resistance and tolerance mechanisms. While the presence of *mecA* remains a reliable indicator of oxacillin resistance, our findings point to a context-dependent, potentially epistatic interaction of chromosomal (tolerance) mutations with *mecA* that shapes the phenotypic outcome. These findings emphasize the need to view resistance and tolerance as genetically interlinked but phenotypically separate survival strategies, an important distinction which challenges diagnostic approaches and treatment decisions based solely on MIC thresholds but has the potential to enhance clinical treatment of infections caused by *S. epidermidis*.

## Materials and Methods

### *S. epidermidis* isolate collection and routine growth conditions

A total of 88 primary *S. epidermidis* isolates were used in this study, of which 80 have previously been described^13,40^. Type strain NCTC11047 (ATCC14990; CCM2124) was purchased from the UK Health and Security Agency. Genome assemblies of all isolates are available through the PubMLST database^78^. Isolate metadata is summarised in Supplementary File S1. All bacterial strains used in this study were routinely grown in Tryptic Soy Broth (TSB, Difco), at 37°C with agitation (180 rpm). For growth on solid media, TSB supplemented with 1.5% (w/v) of bacteriological agar, or Columbia agar plates with 5% defibrinated horse blood (E&O Laboratories) were used, followed by static incubation at 37°C. For long-term storage at -70°C, *S. epidermidis* liquid cultures were supplemented with glycerol to a final concentration of 20% (v/v).

### Phylogenetic analysis

A maximum-likelihood phylogeny was constructed for the 88 *S. epidermidis* strains with tested antibiotic resistance phenotypes. Genome assemblies were annotated using Prokka (v1.14.5)^79^ with the compliant flag. Core-gene alignments were constructed by Panaroo (v1.3.2)^80^ with the remove invalid genes flag and strict stringency filtering. Phylogenies were inferred from core-gene alignments using RAxML (v8.2.12)^81^ with GTR-GAMMA as a substitution model. Recombination was accounted for by ClonalFrameML (v1.13)^82^ using the default settings. The resulting phylogeny was visualised in iTOL (v6.9.1)^83^. For wider contextualisation of the 88 isolates, all other available *S. epidermidis* genomes from PubMLST^78^ were downloaded (n = 1,098, accessed October 2024). For quality control, assemblies that were not 2-3 Mb in length (n = 26) and contained over 357 contigs (n = 160) were removed from further analysis and a phylogenetic tree produced as described above (Supplementary File S1).

### Minimum inhibitory concentration determination

Minimum inhibitory concentrations (MICs) were determined by microbroth dilutions according to the European Committee on Antimicrobial Susceptibility Testing (EUCAST)^84^ guidelines. 10 antibiotics with distinct mechanisms of action regularly used to treat staphylococcal infections were tested. Solvent and range tested were adjusted for each antibiotic as indicated in brackets: oxacillin (0.0625 - 1024 mg L^-1^, dH_2_O, Sigma Aldrich), ciprofloxacin (0.03125 - 128 mg L^-1^, 50 mM acetic acid, Sigma Aldrich), gentamicin (≤0.125-256 mg L^-1^, dH_2_O, Sigma Aldrich), erythromycin (≤0.125 - >1024 mg L^-1^, 70% EtOH, Sigma Aldrich), chloramphenicol (≤0.125 - 128 mg L^-1^, 70% EtOH, Sigma Aldrich), rifampicin (0.00045 - 256 mg L^-1^, DMSO, Sigma Aldrich), trimethoprim (≤0.125 - 128 mg L^-1^, DMSO, MPBiomedicals), tetracycline hydrochloride (≤0.125 - 128 mg L^-1^, dH_2_O, Cayman Chemical), vancomycin hydrochloride (≤0.125 - 128 mg L^-1^, dH_2_O, Cayman Chemical), linezolid (≤0.125-128 mg L^-1^, DMSO, Thermo Scientific). Briefly, antibiotics at 2x maximum concentration were 2-fold serially diluted across 96-well plates in Muller-Hinton II broth (MHB, Thermo Fisher Scientific). Plates were inoculated with a final concentration of approximately 5x10^5^ colony forming units (CFU) mL^-1^ and incubated for 20-24 hours at 37°C before reading absorbance at optical density of 600 nm (OD_600_). Absorbance was normalised against untreated control wells before calling MIC_90_, where growth was ≥0.1 OD_600_. Strains were classified as sensitive or resistant using EUCAST breakpoints (v. 14.0)^84^. An overview of MICs is given in Supplementary File S2.

### Detection of acquired and chromosomal antibiotic resistance determinants

Whole-genome sequences of the 88 isolates analysed in this study were obtained from the *S. epidermidis* genome collection in PubMLST^78^ and screened for known genomic antimicrobial resistance determinants. Briefly, a query file containing the protein sequences of all resistance determinants of interest was compiled from different sources, including the CARD database^59^ and relevant literature^17,22,85,86^. We then performed a BLAST search (tBLASTn, BLAST+ v2.16.0)^87^ to identify homologous hits between the query proteins and each isolate assembly. Any hits with <70% identity, or which overlapped another hit with a higher bit-score, were excluded. The remaining hits were translated to protein using Transeq (EMBOSS v6.6.0)^88^ and aligned to their corresponding original queries using MAFFT (v7.526)^89^ to identify putative frameshifts, premature stop codons, and to calculate the percentage identity and coverage. Hits with less than 85% identity at 100% coverage, and genes with premature stop codons and frameshifts were termed ‘absent’ unless otherwise indicated (e.g. partial alleles of *mecR* and *mecI* genes).

For resistance caused by chromosomal variation in core genes rather than acquired genes, amino acid substitutions and insertions/deletions were identified by comparison to the NCTC11047/ATCC14990 reference genome. To achieve this, the previously filtered nucleotide sequences were extracted from the sample assemblies. Hits in different reading frames were merged using a maximum window of 15 bp to obtain a single contiguous nucleotide sequence. Where there was more than one hit per gene due to genes spanning contig boundaries, a single consensus sequence was generated, where overlap discrepancies were encoded with ‘N’. The resulting nucleotide sequences for all samples were combined into a single nucleotide FASTA file (one per gene) and aligned using MAFFT (v7.526)^89^. A custom Python script (https://github.com/Sheppard-Lab/AlignmentToVariants) was then used to identify variants with NCTC11047/ATCC14990 as the nucleotide reference, which were compared to resistance mutations described in databases and the literature^59^. For trimethoprim resistance mediated by Dfr, manual analysis was performed based on the PubMLST^78^ entries. The full list of resistance determinants with database accessions can be found in Supplementary file S3.

### Determination of Staphylococcal Cassette Chromosome *mec* type

To determine the Staphylococcal Cassette Chromosome *mec* (SCC*mec*) type causing β-lactam resistance, we developed a Linux-based custom python script of SCC*mec* classifier. The complete source code for this tool is available on GitHub (https://github.com/Sheppard-Lab/SCCmecClassifier). Briefly, by integrating the alignment tool Minimap2^90^ with a previously available web-based SCC*mec*Finder tool^91^, we developed a new Linux-based SCC*mec* classifier to handle multiple assemblies concurrently with a reduction in processing time.

The gene database for this tool was compiled following the guidelines detailed by a recent review^19^ and contains all types of *mec* gene complexes and *ccr* gene complexes from each SCC*mec* type in the reference genome. Using Minimap2^90^ with a divergence level set at 5%, sequence alignment map (SAM) and browser extensible data (BED) files, which contain alignment statistics, were generated. From these SAM and BED files, the tool filters the best matching *mec* gene, *mec* complex and *ccr* complex types. Given the potential for multiple *ccr* types within the genome, if two best matches were within a 10% difference, the *ccr* gene closest to the *mecA* or *mecC* gene was prioritized. Considering that assemblies are composed of multiple contigs, there are risks that genes located at contig ends lead to incorrect alignment results. To address this, the program iterates up to three times to identify hidden matches in other contigs. The output includes a statistics file that lists the matching percentage and the contigs where the genes are located (Supplementary file S4).

### Directed evolution of *mecA*^-^ and *mecA*^+^ isolates for increased oxacillin tolerance and resistance

Tolerance Disk Tests (TDtests)^92,93^ were used to select a subset of *mecA*^-^ (n = 8) and *mecA*^+^ (n = 16) isolates for increased oxacillin tolerance and resistance. Overnight cultures were diluted to a final OD_600_ of 0.5 in phosphate buffered saline (PBS) and streaked with a swab on tryptic soy agar (TSA) plates. Antibiotic disks with 1 µg or 10 µg oxacillin for *mecA*^-^ and *mecA*^+^ isolates, respectively, were placed on the plates, with blank disks used as negative control.

After incubation at 37°C for 18–20 h, antibiotic disks were replaced with disks soaked with 10 μL of 40% glucose and 20% casamino acids. For each TDtest, three colonies growing within the inhibition zone were picked after 24 h incubation at 37°C and streaked to single colony. Modal oxacillin MIC values (most frequently repeated MIC value) for evolved isolates were determined by broth microdilution as described above, using the ancestral isolates as reference. Information on all strains that have undergone oxacillin selection, and their ancestral isolates are summarised in Table S1 and S2, and Supplementary File 6.

### Sequencing of adapted *mecA*^-^ and *mecA*^+^ isolates and variant calling

Genomic DNA of 96 ancestral and evolved *mecA*^-^ and *mecA*^+^ isolates was extracted using the Maxwell RSC Cultured Cell DNA Kit (Promega) on semiautomated DNA extraction machines (Maxwell RSC Instrument, Promega), following the manufacturer’s instructions. To facilitate cell lysis, samples were treated with 0.125 mg mL^-1^ lysostaphin (Sigma Aldrich, ≥3000 unit mg^-1^) dissolved in 20 mM NaOAc (pH 5.2) at 37°C for 1 – 2 h at 180 rpm. DNA was quantified using the QuantiFluor ONE dsDNA System and a Quantus Fluorometer (Promega) according to manufacturer’s instructions. Libraries were prepared using the Illumina® DNA Prep, (M) Tagmentation kit with Unique Dual Index adapters, generating an insert size of approximately 350 bp. Pooled libraries were loaded at 750 pM onto an Illumina® NextSeq™ 2000 using XLEAP-SBS™ P2 reagents for a paired-end 300-cycle run (2 x 150 bp).

The resulting short reads were assembled using shovill (v1.1.0) using the trim flag to remove any remaining adapter sequences (https://github.com/tseemann/shovill). Whole-genome sequencing datasets of evolved isolates were deposited in the Sequence Read Archive (SRA) operated by the National Center for Biotechnology Information (NCBI) under BioProject accession number: PRJNA1314787. SRA accession numbers are given in Supplementary File S1. Assemblies were annotated using Prokka (v1.14.5)^79^ using the compliant flag. Genomic variation in evolved strains was identified using snippy (v4.6.0, https://github.com/tseemann/snippy) using the respective ancestral strain as reference. Amino acid altering (missense) or aborting (non-sense) mutations, as well as insertions and deletions are given in Table S1 and S2. A summary of all mutations including synonymous nucleotide changes are given in Supplementary File S5. Raw sequencing reads of evolved isolates were deposited in the Sequence Read Archive (SRA) operated by the National Center for Biotechnology Information (NCBI) under BioProject accession number: PRJNA1314787. ChimeraX (v1.8) was used to visualise the 11 amino acid changes in GdpP, that were observed in the directed evolution experiments^94^.

### Time-kill curves to measure antibiotic tolerance

Time-kill assays were performed for oxacillin to test for increased tolerance. Overnight cultures were diluted to an OD_600_ of 0.02 in 6 mL of TSB and exposed to oxacillin at 100-fold modal MIC (6.25 - 25 mg L^-1^). 1 mL samples were taken immediately after the antibiotic challenge (t_0_ = 0 h), and cultures further incubated at 37°C with continuous shaking (180 rpm) and 1 mL samples were taken in 2 h intervals for 6 h (t1−3). Samples were centrifuged at 17,000×*g* for 2 min and washed twice with 1 mL Dulbecco’s Phosphate Buffered Saline (PBS, Sigma-Aldrich) to remove residual antibiotic. Samples were 5-fold serially diluted in PBS and 10 μL were spotted on TSA plates using liquid handler OT-2 (Opentrons). Plates were incubated at 37°C static incubation and colonies were counted after 24 h to determine colony-forming units (CFU) ml^-1^. Relative survival was quantified by dividing CFU ml^-1^ at each time by CFU ml^-1^ at t_0_, resulting in 100% relative survival at t_0_. Experiments were conducted in at least 3 biological replicates. The minimum duration to kill 99% of the bacterial population was calculated by extrapolating the missing value from the standard curve^95^.

### Growth curves

Growth kinetics of *S. epidermidis* isolates were acquired using a VANTAstar (BMG Labtech) multimode plate reader. *S. epidermidis* cultures were diluted 1:200 from overnight cultures into fresh TSB medium in 96 well plates. The growth dynamics were measured every 5 min using a 2x2 matrix scan (2 mm diameter) and 500 rpm double orbital shaking at 37°C for 20 h. For each isolate, at least three technical repeats were averaged, with at least 4 biological repeats overall. Representative growth curves are displayed.

### Population genomics analyses

Genomic variation among isolates from directed evolution experiments was contextualised by comparison to 1000 *S. epidermidis* isolate genomes available in the PubMLST database^78^ using the GeneScanner plugin^96^ (https://github.com/Sheppard-Lab/GeneScanner). Nucleotide sequences of 11 genes that had two or more independent mutations in the directed evolution experiments were used to query against isolate assemblies using blastn (BLAST 2.12.0+ with settings: word size: 20; reward: 2; penalty: -3; gapopen: 5; gapextend: 2)^87^. The identified matching sequences were extracted and aligned using MAFFT (v7.505) with default parameters to create the nucleotide alignment input file^89^. GeneScanner then identifies the frequency of synonymous and non-synonymous mutations, stop codons, insertions, and deletions across aligned sequences. It applies two quality filters relative to a reference sequence, requiring at least 80% ungapped coverage and 80% pairwise identity. In nucleotide analysis mode, the program scans each alignment codon by codon while maintaining the correct reading frame, comparing translated codons to the reference to distinguish synonymous from non-synonymous changes. All mutation data are then compiled into a spreadsheet. The identified sequence variation was then manually compared to mutational sites reported in the evolution datasets.

### Statistical tests

All statistical analyses were conducted using GraphPad Prism (v10.1.2). Fisher’s exact test was used to infer the association between resistance determinant and drug phenotype. Mann Whitney U test was used to analyse the changes in MIC. Threshold for significance was set at *P* < 0.05 for all tests.

## Abbreviations

AMR: Antimicrobial resistance
MIC: Minimum inhibitory concentration
SCC*mec*: Staphylococcal Cassette Chromosome *mec*
ST: sequence type
MDK_99_: Minimum duration to kill 99% of the population
OXA: Oxacillin
CIP: ciprofloxacin
GEN: gentamicin
ERY: erythromycin
CHL: chloramphenicol
RIF: rifampicin
TMP: trimethoprim
TET: tetracycline
VAN: vancomycin
LNZ: linezolid
MDR: Multidrug resistance
TSA/TSB: Tryptic Soy Agar/Broth
SAM: sequence alignment map
BED: browser extensible data.

## Declarations

### Ethics approval and consent to participate

Not applicable.

### Consent for publication

Not applicable.

### Availability of data and materials

Whole-genome sequencing datasets of evolved isolates were deposited in the Sequence Read Archive (SRA) operated by the National Center for Biotechnology Information (NCBI) under BioProject accession number: PRJNA1314787. Genome assemblies of primary isolates are available on PubMLST. Isolate IDs, MIC values, SCC*mec* analysis output, and variant analysis of evolved isolates can be found in the supplementary files.

AMR variants were identified using a custom python script (https://github.com/Sheppard-Lab/AlignmentToVariants). Code for running *SCCmec* classification is available under https://github.com/Sheppard-Lab/SCCmecClassifier. GeneScanner code and detailed requirements for each release version are publicly available (https://github.com/Sheppard-Lab/GeneScanner, DOI: 10.5281/zenodo.17495646.

### Authors’ contributions

Conceptualization of project: CMK, SKS, MS

Methodology: CMK, JCL, EAC, SK, PSR, KAJ, SKS, MS

Data acquisition: CMK, CLM, JCL, EAC, CW, SK

Funding acquisition: CMK, SKS, MS

Project administration and supervision: CMK, SKS, MS

Writing - original draft: CMK, MS, SKS

Writing - review & editing: all authors.

## Acknowledgements

We thank David Baker (Quadram Institute) for his generous support and technical expertise in sequencing the isolates.

## Figures

**Figure S1:**
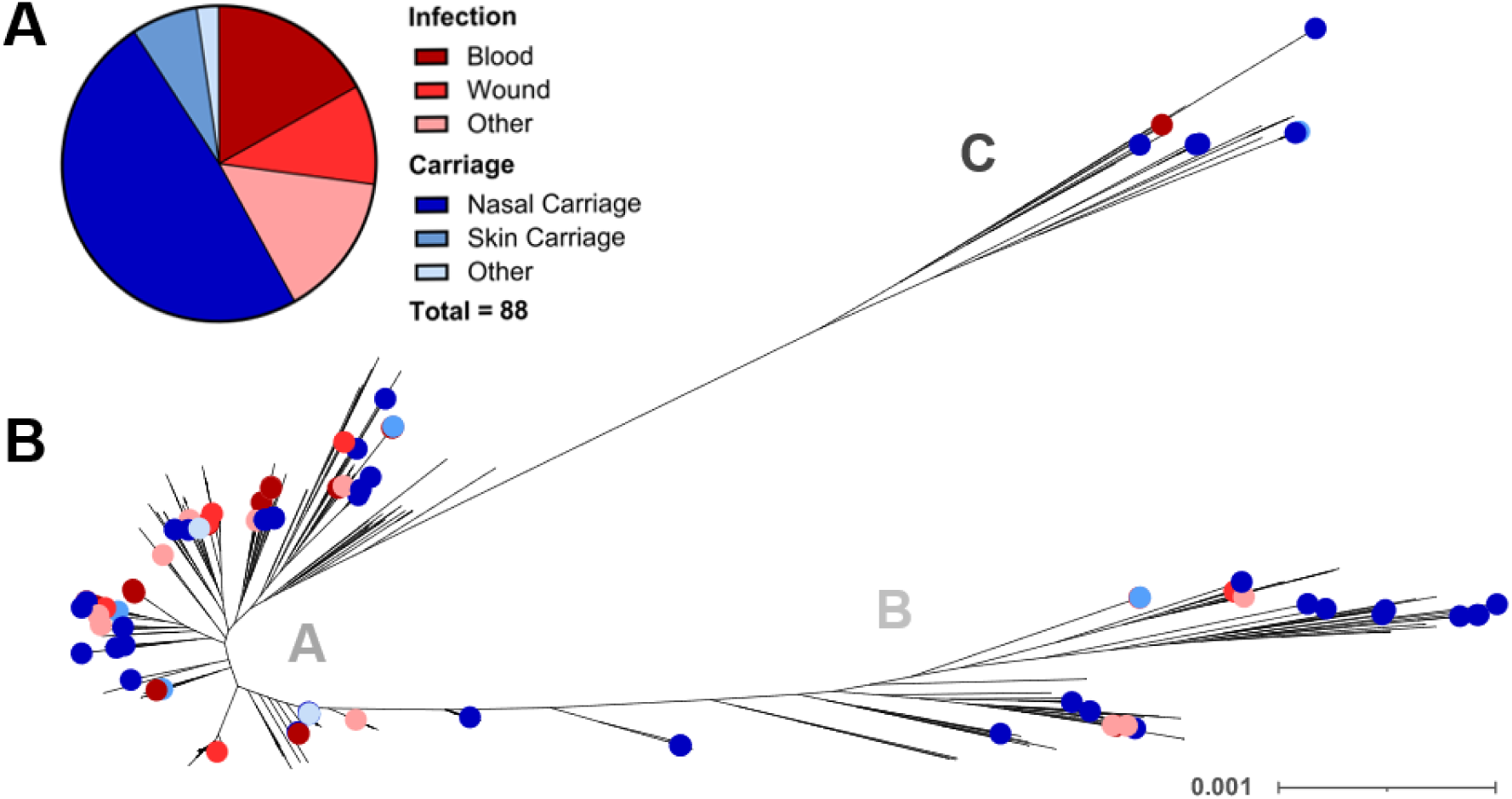
Topology of *S. epidermidis* collection is consistent with population structure of other *S. epidermidis* isolates. Isolation source of 88 *S. epidermidis* infection (n = 37) and carriage (n = 51) isolates used in this study. Shades of red correspond to infection sources, shades of blue to isolation from asymptomatic carriage. B: Maximum likelihood phylogenetic tree of 1000 *S. epidermidis* isolates, from a core genome alignment of all genes shared by > 95% isolates (n = 1946), with isolation source of 88 isolates used in this study highlighted in colour, corresponding to those used in panel A. The tree scale represents the number of substitutions per site. Previously reported distinct clades A, B, and C are indicated in grey letters.

**Figure S2:**
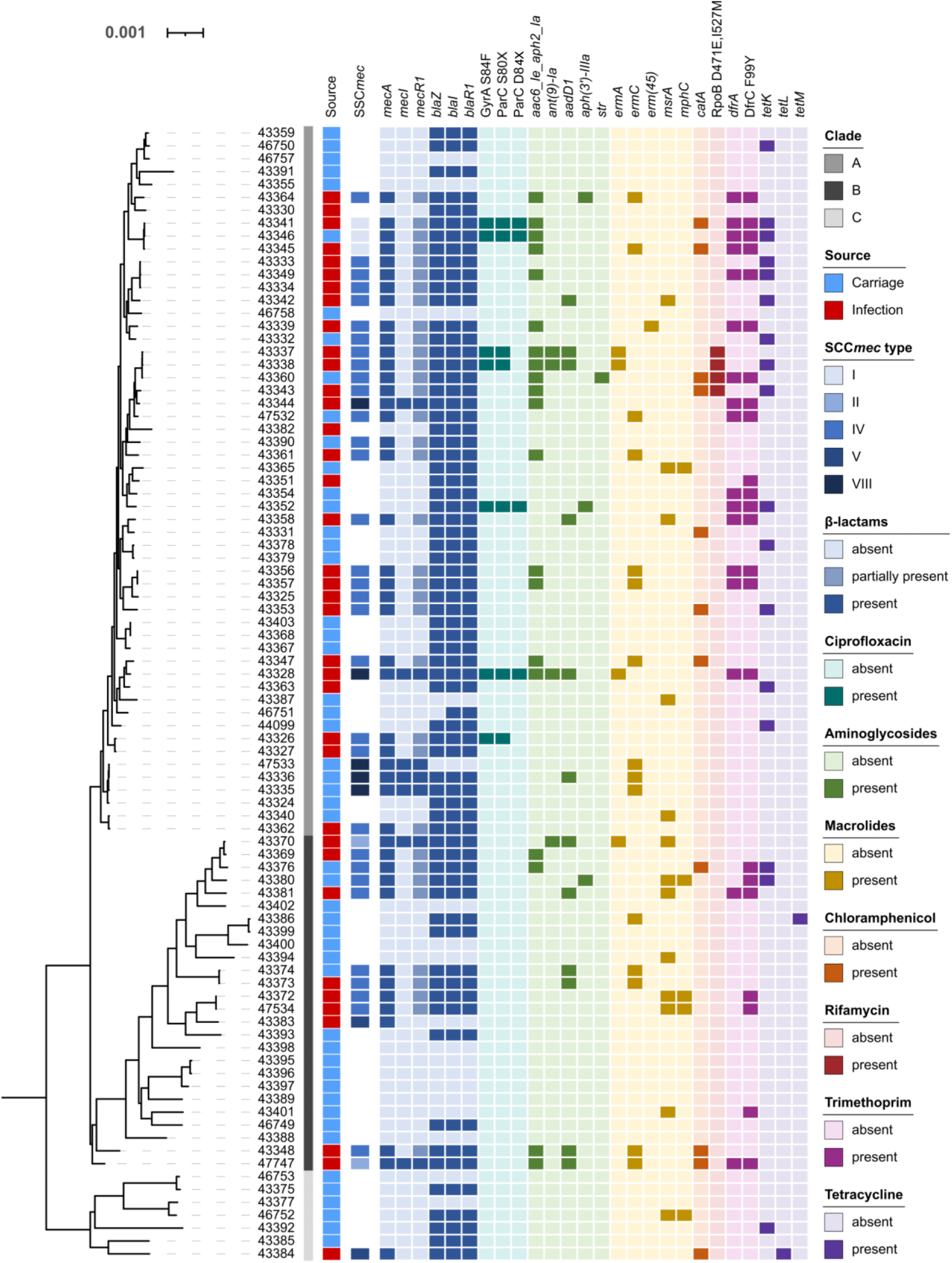
Phylogenetic distribution of genotypic antibiotic resistance determinants matches phenotypically multidrug resistant *S. epidermidis* clusters. Presence and absence of AMR genes, and amino acid changes, that are associated with AMR to the eight antibiotics tested are displayed alongside their phylogeny. The tree scale represents the number of substitutions per site. Different shades of grey and red or blue indicating placement in deep branching clades A-C, and isolation from infections or carriage, respectively. Presence of an acquired AMR gene and amino acid changes are indicated through dark shades of the respective colour, while absence thereof corresponds to the lightest shade. β-lactam (dark blue), fluoroquinolone (teal), aminoglycoside (green), macrolide (yellow), phenicol (orange), rifamycin (dark red), trimethoprim (pink), tetracycline (purple). For β-lactam resistant isolates, the SCC*mec* type is given in shades of blue. Further, partial presence of *mecR1* as part of specific SCC*mec* cassettes is indicated in lighter blue shading.

**Table S1:**
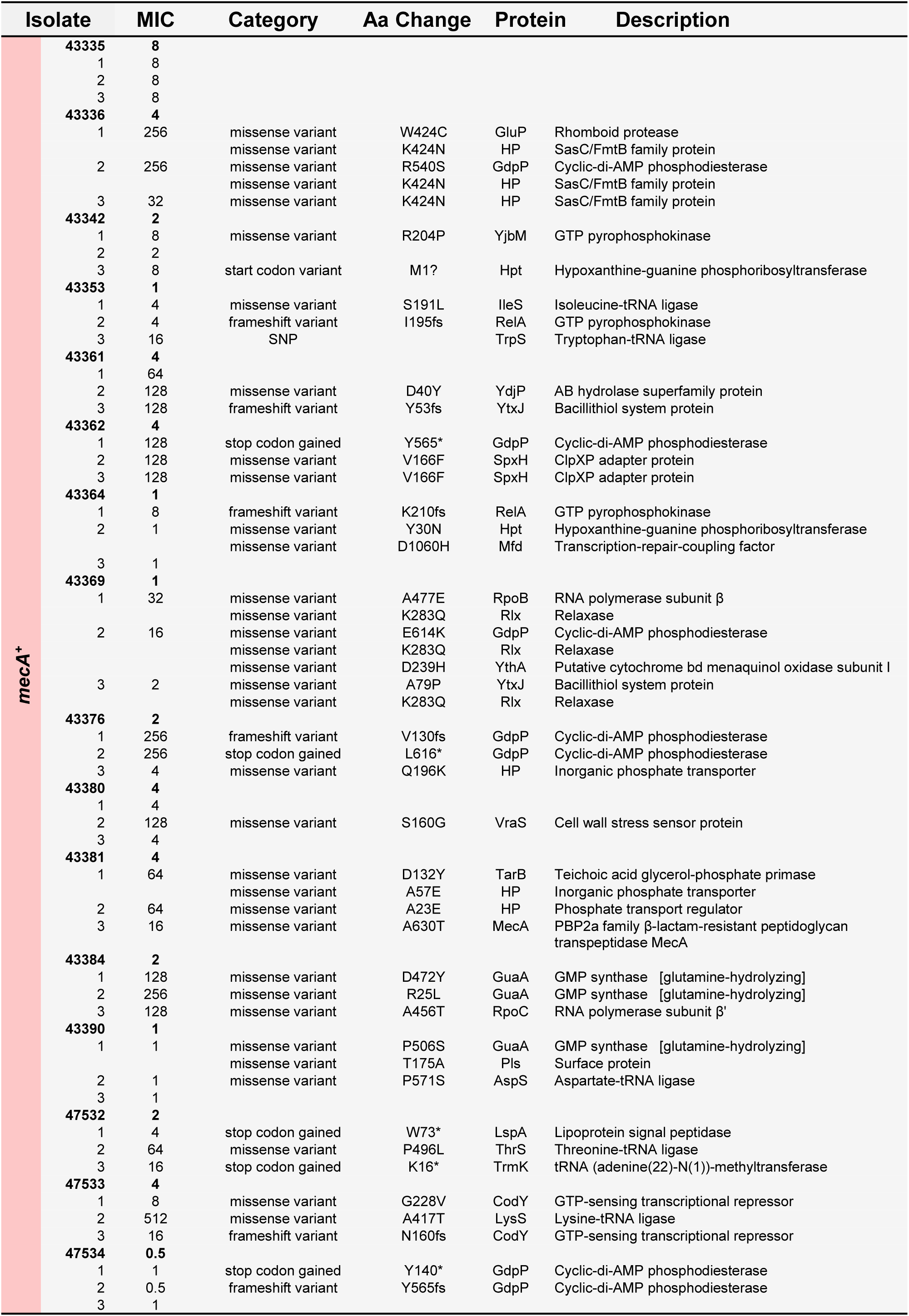
Non-synonymous mutations identified in *mecA^+^* isolates under antibiotic pressure. Isolate ID is given with 1, 2, and 3 indicating three evolved isolates. Oxacillin MICs are given in mg L^-1^. Single letter code is used to refer to amino acids, * stands for non-sense variants (stop codon) and fs for frame shift mutations.

**Figure S3:**
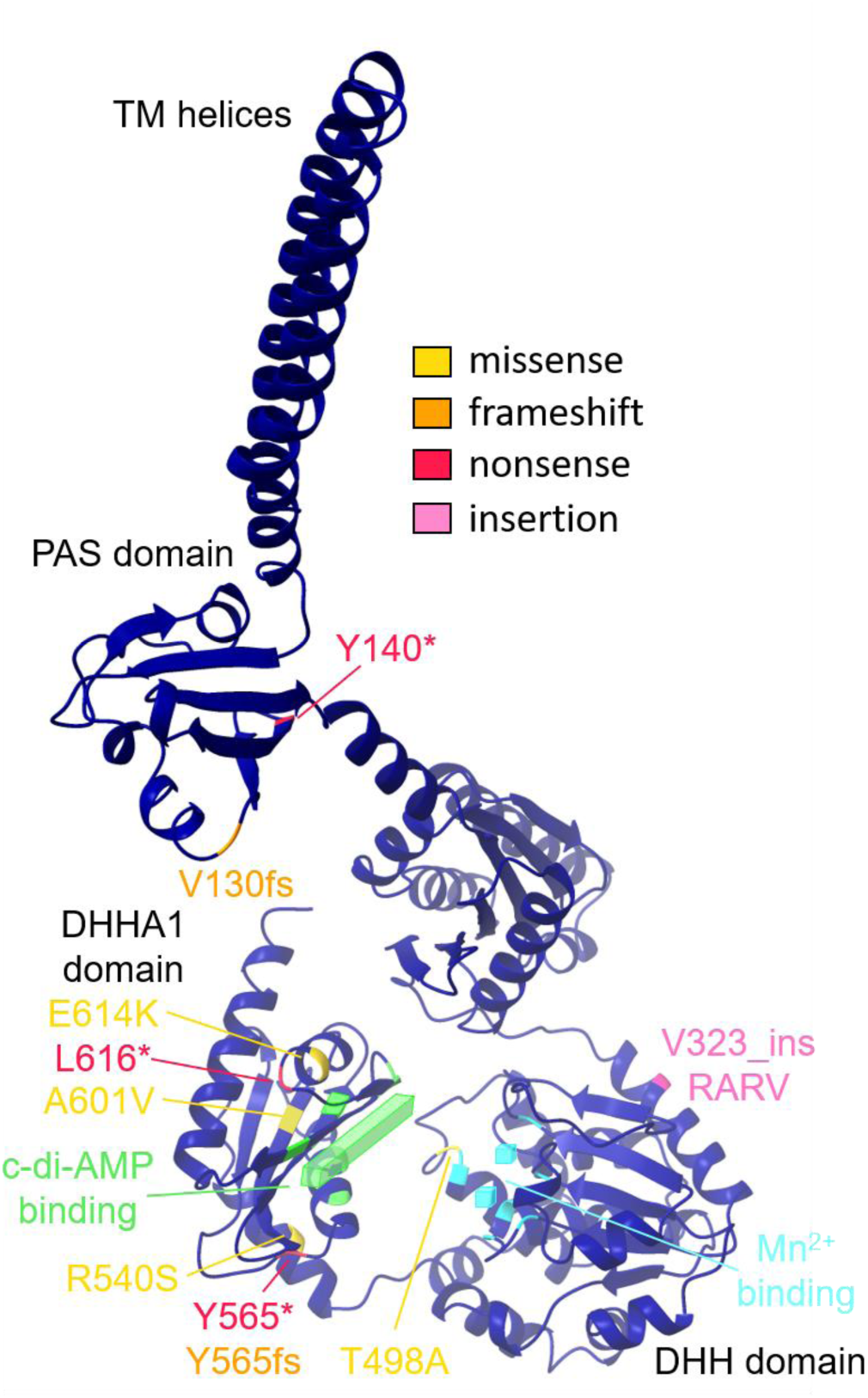
Amino acid changes indicate a loss of GdpP function. The majority of the 11 non-synonymous changes observed across directed evolution dataset in the cyclic di-AMP phosphodiesterase GdpP accumulate around the substrate (c-di-AMP) and cofactor (Mn2^+^) binding sites, indicting a potential reduction or loss of GdpP function.

**Table S2:**
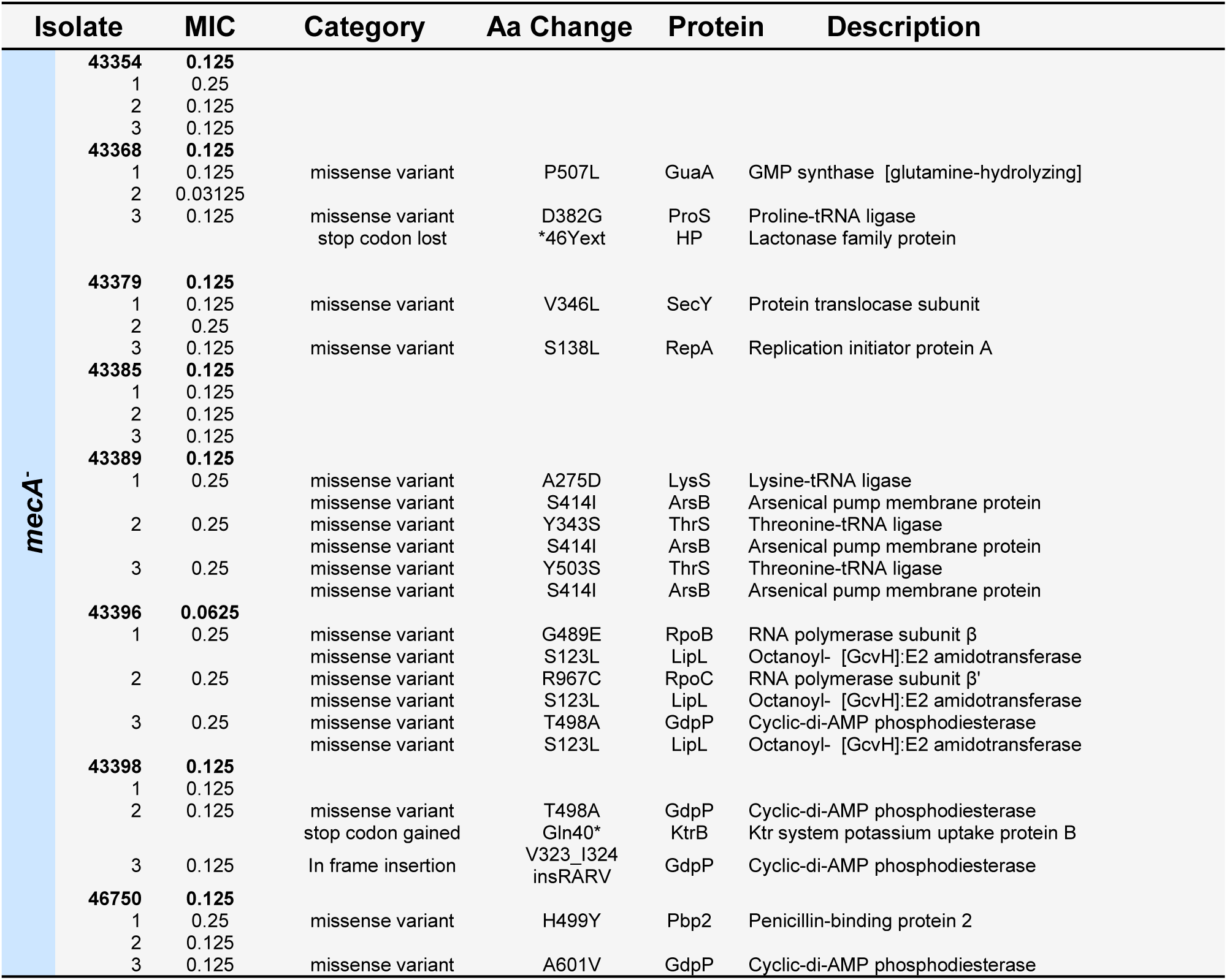
Non-synonymous mutations identified in *mecA^-^* isolates under antibiotic pressure. Isolate ID is given with 1, 2, and 3 indicating three evolved isolates. Oxacillin MICs are given in mg L^-1^. Single letter code is used to refer to amino acids, * stands for non-sense variants (stop codon) and fs for frame shift mutations.

**Figure S4:**
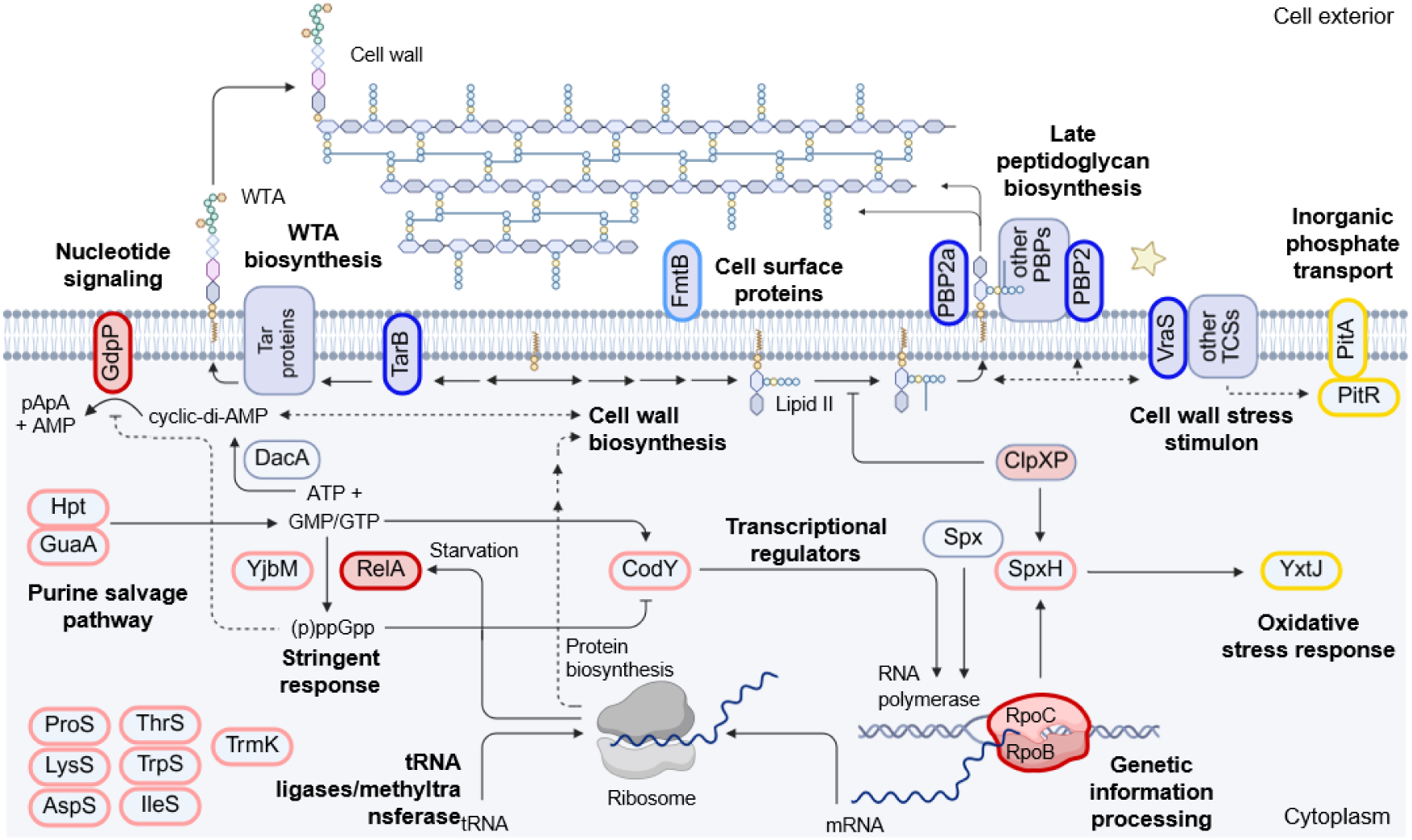
Isolates adapted to oxacillin have changes in auxiliary and tolerance/potentiating factors. Schematic of core processes of the cell, with all proteins highlighted that had protein-altering changes in the directed evolution assay. Proteins targeted by non-synonymous changes can be broadly grouped into two groups: potentiator/tolerance genes (red) and auxiliary factors (blue). Red fill: protein has previously been named a ‘potentiator’ to β-lactam resistance in *S. aureus*. Red outline: Protein-altering changes were observed and the gene previously linked to tolerance. Pink outline: Amino acid change identified in this study. Blue outline: Protein names an auxiliary factor with amino acid changes observed in this study. Light Blue: amino acid changes in protein of same family was identified in this study. Yellow: Proteins in other pathways that had two or more independent amino acid changes. Created in BioRender. Kobras, C. (2026).

**Figure S5:**
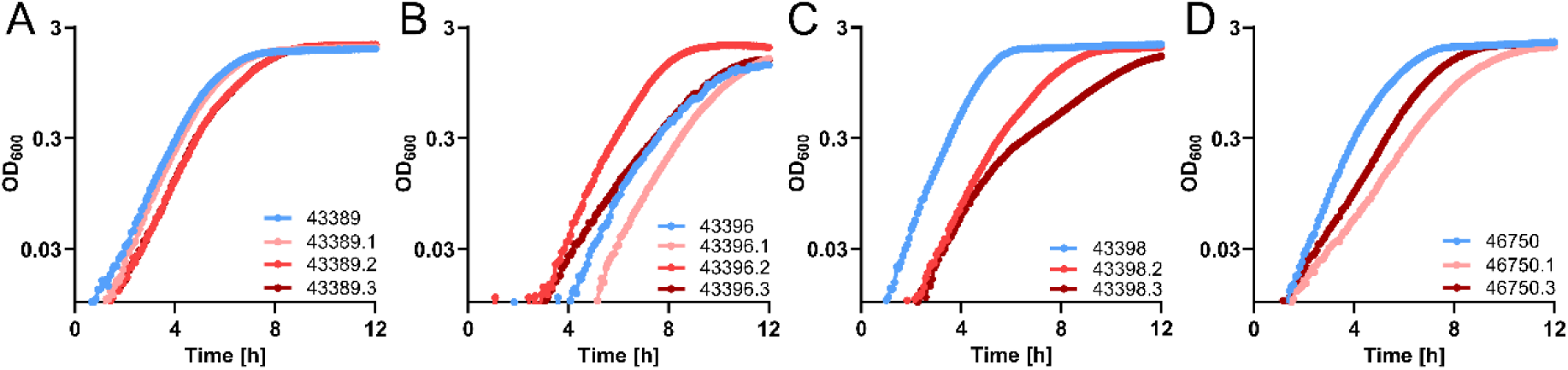
Several evolved isolates, but not all, display altered growth behaviour and slower growth. Growth curves were assessed for four ancestral strains and two or three corresponding evolved isolates carrying non-synonymous mutations. Evolved isolates of strain 43389 carrying mutations in tRNA ligases did not show altered growth despite increased antibiotic survival (A), while several oxacillin-tolerant isolates exhibited reduced growth, consistent with growth reduction as a common tolerance mechanism (B–D). All growth curves are representative of at least four biological replicates.

**Figure S6.**
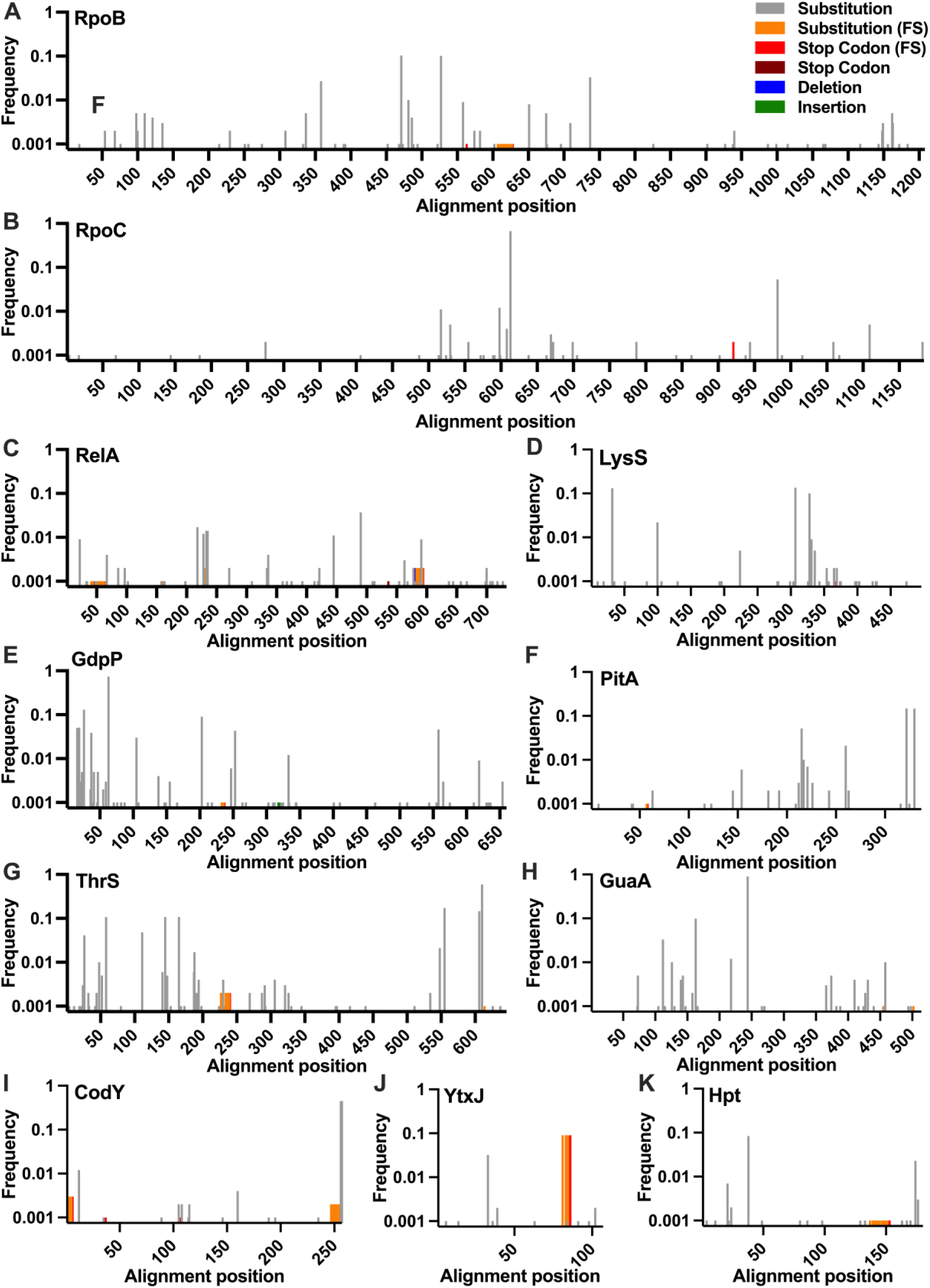
Genetic variation is prevalent across natural *S. epidermidis* isolate genomes. GeneScanner analysis of 1,000 natural *S. epidermidis* genomes identified genetic variation in genes that acquired two or more nonsynonymous mutations during the evolution experiment, compared to ATCC 14990 reference alleles. Frequencies are given for amino acid substitutions in grey, substitutions following a frameshift (FS) in orange, stop codons following frameshifts (FS) in red, stop codons through non-synonymous substitutions in dark red, in-frame amino acid deletions in blue, and in-frame insertions in green. A: RpoB, B: RpoC, C: RelA, D:, LysS, E: GdpP, F: PitA, G: ThrS, H: GuaA, I: CodY, J: YtxJ, K: Hpt.

